# Zika virus infection in Collaborative Cross mice

**DOI:** 10.1101/695510

**Authors:** Melissa D. Mattocks, Kenneth S. Plante, Ethan J. Fritch, Ralph S. Baric, Martin T. Ferris, Mark T. Heise, Helen M. Lazear

**Affiliations:** Departments of Microbiology & Immunology, University of North Carolina at Chapel Hill; Departments of Genetics, University of North Carolina at Chapel Hill; Departments of Epidemiology, University of North Carolina at Chapel Hill

## Abstract

The 2015-2016 emergence of Zika virus (ZIKV) in the Americas, and recognition that ZIKV infection during pregnancy can result in birth defects, revealed a need for small animal models to study ZIKV pathogenic mechanisms and evaluate candidate vaccines and antivirals. Mice would be an attractive system for such studies, but ZIKV replicates poorly in laboratory mice because it fails to antagonize murine STAT2 and STING. To address this, most ZIKV pathogenesis studies have used mice with impaired interferon signaling (e.g. *Ifnar1*^−/−^ or treatment with IFNAR1-blocking antibodies). However, using mice with severe defects in innate antiviral signaling confounds studies of viral pathogenic mechanisms. Collaborative Cross (CC) mice have proven to be a valuable system for developing new mouse pathogenesis models for viral infections that are not well modeled in conventional laboratory mouse lines. To test whether CC mice could provide an immune-competent model for ZIKV pathogenesis, we infected CC lines with ZIKV and assessed weight loss, viremia, and production of neutralizing antibodies. We tested 21 CC lines (CC001, CC002, CC003, CC004, CC005, CC006, CC011, CC012, CC013, CC019, CC024, CC028, CC040, CC041, CC042, CC046, CC051, CC059, CC061, CC068, and CC072, 13 of which have non-functional alleles of the flavivirus restriction factor Oas1b) and 3 ZIKV strains (MR766, H/PF/2013, and a mouse-adapted variant of Dakar 41525). ZIKV infection did not induce weight loss compared to mock-infected controls and accordingly only low levels of viral RNA were detected in serum. Only a subset of mice developed neutralizing antibodies to ZIKV, likely due to overall low levels of infection and viremia. Our results are consistent with other studies demonstrating poor ZIKV infection in interferon-intact mice and suggest that the tested CC lines do not include polymorphic host genes that greatly increase susceptibility to ZIKV infection.

## Introduction

The 2015-2016 emergence of Zika virus (ZIKV) in the Americas, and recognition that ZIKV infection during pregnancy can result in birth defects, revealed a need for small animal models to study ZIKV pathogenic mechanisms and evaluate candidate vaccines and antivirals (1). Mice would be an attractive system for such studies due their low cost and genetic tractability and mice have proven to be a valuable system for studying other flaviviruses, such as West Nile virus (WNV). However, studies of ZIKV in mice are confounded because mouse type I interferon (IFN-αβ) signaling restricts ZIKV replication (2), in part due to the inability of ZIKV to antagonize murine STAT2 and STING (3-6). This results in diminished ZIKV replication in immune competent mice and has led most groups to use IFN-deficient mouse models including mice lacking the type I and/or type II IFN receptors (e.g., *Ifnar1*^−/−^ or *Ifnar1*^−/−^ *Ifngr1*^−/−^ double knockout), mice with defects in IFN induction or signaling (e.g., *Irf3*^−/−^ *Irf5*^−/−^ *Irf7*^−/−^ or *Stat2*^−/−^), or mice treated with IFNAR1-blocking antibody (7). These models are useful because they allow robust ZIKV replication and produce severe disease outcomes with high penetrance. However, these models have significant limitations for studying immune mechanisms that control ZIKV replication and disease. In particular, IFN signaling plays a key role in restricting flavivirus tropism and IFN-deficient mice typically develop disseminated ZIKV infection that does not recapitulate the tissue tropism observed in humans. Thus, there is a need for new mouse models that exhibit robust ZIKV replication and disease within an immune competent host.

The Collaborative Cross (CC) is a mouse genetic reference population that was developed to study complex trait genetics and to model genetically complex populations, such as humans (8, 9). CC mice are derived from eight founder lines: five founder lines are classical laboratory lines with an extensive history of use in mouse genetics, microbial pathogenesis, and disease models (C57BL/6J, 129S1/SvImJ, A/J, NOD/ShiLtJ, and NZO/HiLtJ). The other three CC founder lines are wild-derived inbred lines, which represent the three major subspecies of Mus musculus: casteneus (CAST/EiJ), musculus (PWK/PhJ), and domesticus (WSB/EiJ), and introduce much of the genetic diversity into the CC. These eight founder strains were interbred in a funnel breeding scheme to produce recombinant mice with genomic contributions from each founder, which were then bred to homozygosity to produce approximately 80 recombinant inbred CC lines. The CC mouse genetic reference population is designed to model the genetic diversity present in human populations and allow the identification and study of polymorphic host genes underlying complex phenotypes, including the immune response to viral infection. Because each CC line is inbred, they provide a reproducible system to study diverse phenotypes.

The genetic architecture of the CC results in a broad spectrum of antiviral responses, including responses not observed in standard mouse lines that more closely resemble the clinical outcomes observed in humans (8, 9). This has resulted in the development of mouse models for pathogens, such as Ebola virus, severe acute respiratory syndrome (SARS) coronavirus, WNV, and chikungunya virus, that better model clinical aspects of human disease compared to standard laboratory mouse lines (10-12). Therefore, we sought to use the CC to identify polymorphic host genes that control ZIKV specific innate and adaptive immunity and determine whether variation in these genes impacts ZIKV susceptibility. These experiments would have the potential to identify CC lines with enhanced susceptibility to ZIKV replication and disease, thereby resulting in the development of new mouse models of ZIKV pathogenesis.

## Methods

### Mice

CC mice were obtained from the UNC Systems Genetics Core Facility. All mouse procedures were performed under protocols approved by the UNC Insitutional Animal Care and Use Committee. 5-week-old male and female mice or 15-21-week-old male mice were inoculated in a volume of 50µL by a subcutaneous (footpad) route. Mice received 1 × 10^3^ FFU of ZIKV strain H/PF/2013 or MR766 or 1 × 10^5^ FFU of strain Dakar-MA; mock-infected mice received diluent (HBSS with Ca^2+^ and Mg^2+^ supplement with 1% heat-inactivated fetal bovine serum (FBS)). Mice were weighed and observed for disease signs daily for 14 days, at which time tissues were harvested following perfusion with PBS. In separate experiments to evaluate viremia, blood was collected at 1, 3, and 5 days post-infection (dpi) by submandibular bleed with a 5mm Goldenrod lancet.

### Cells and viruses

Vero (African green monkey kidney epithelial) cells were maintained in Dulbecco’s modified Eagle medium (DMEM) containing 5% heat-inactivated FBS at 37°C with 5% CO_2_. ZIKV strain H/PF/2013 was provided by the U.S. Centers for Disease Control and Prevention (13). ZIKV strain MR766 was obtained from the World Reference Center for Emerging Viruses and Arboviruses; the MR766 variant used in these studies lacks a glycosylation site in the viral E protein (14-17). ZIKV strain Dakar-MA was generated by serially passaging strain Dakar 41525 in *Rag1*^−/−^ mice to generate a mouse-adapted variant and was provided by Dr. Michael Diamond (Washington University in St. Louis) (4). Virus stocks were grown in Vero cells and titered by focus-forming assay (FFA) (18). Duplicates of serial 10-fold dilutions of virus in viral growth medium (DMEM containing 2% FBS and 20 mM HEPES) were applied to Vero cells in 96-well plates and incubated at 37°C with 5% CO_2_ for 1 hr. Cells were then overlaid with 1% methylcellulose in minimum essential medium Eagle (MEM). Infected cell foci were detected 42-46 hpi. Following fixation with 2% paraformaldehyde for 1 hr at room temperate, plates were incubated with 500 ng/ml of flavivirus cross-reactive mouse MAb E60 (19) for 2 hr at room temperature or overnight at 4°C. After incubation at room temperate for 2 hr with a 1:5,000 dilution of horseradish peroxidase (HRP)-conjugated goat anti-mouse IgG (Sigma), foci were detected by addition of TrueBlue substrate (KPL). Foci were quantified with a CTL Immunospot instrument.

### Measurement of viremia

Blood was collected in serum separator tubes (BD) and serum was separated by centrifugation at 8000rpm for 5 min. Serum was stored at −80°C until RNA isolation. RNA was extracted with the RNeasy Mini Kit (tissues) or Viral RNA Mini Kit (serum) (Qiagen). ZIKV RNA levels were determined by TaqMan one-step quantitative reverse transcription PCR (qRT-PCR) on a CFX96 Touch Real-Time PCR Detection System (BioRad) using standard cycling conditions. Viral burden is expressed on a Log_10_ scale as viral RNA equivalents per ml after comparison with a standard curve produced using serial 10-fold dilutions of RNA extracted from a ZIKV stock. Primers used to detect ZIKV H/PF/2013 were: forward, CCGCTGCCCAACACAAG; reverse, CCACTAACGTTCTTTTGCAGACAT; and probe, /56-FAM/AGCCTACCT/ZEN/TGACAAGCAATCAGACACTCAA/3IABkFQ/ (Integrated DNA Technologies) (20). Primers used to detect ZIKV Dakar-MA were: forward, CCACCAATGTTCTCTTGCAGACATATTG; reverse, TTCGGACAGCCGTTGTCCAACACAAG; and probe: /56-FAM/AGCCTA/ZEN/CCTTGACAAGCAGTC/3IABkFQ (4). Primers and probes were purchased from Integrated DNA Technologies.

### Neutralization assay

Neutralizing antibodies were measured by focus reduction neutralization test (FRNT). Serial 5-fold dilutions of heat-inactivated serum were added to 50-80 focus forming units of ZIKV strain H/PF/2013 or MR766 and incubated for 1 hour at 37°C, then titrated by FFA as above. FRNT50 indicates the serum dilution at which 50% of infectious virus is neutralized. FRNT50 values were calculated with the sigmoidal dose-response (variable slope) curve in Prism 7 (GraphPad), constraining values between 0 and 100% relative infection. A valid FRNT50 curve required an R^2^ >0.75, hill slope absolute value >0.5, and had to reach at least 50% relative infection within the range of the serum dilutions in the assay.

### Data analysis

Data were analyzed with GraphPad Prism software. Weights were compared using two-way ANOVA. For viremia and neutralization analyses, the log-transformed titers were analyzed by the Mann-Whitney test. A p value of < 0.05 was considered statistically significant.

## Results

### Zika virus infection does not induce weight loss in 13 Collaborative Cross lines

To determine whether CC mice could provide a superior pathogenesis model, and potentially to identify polymorphic host genes that contribute to ZIKV pathogenesis, we evaluated ZIKV infection in 21 CC lines (**Table 1**). Because *Oas1b* is well-known as a dominant restriction factor of flavivirus infection in mice, both in conventional laboratory mouse lines and in CC lines (12, 21, 22), we focused our study on lines that carry non-functional *Oas1b* alleles, although some lines with functional *Oas1b* alleles were included for comparison. To determine whether any CC lines were susceptible to ZIKV disease, we infected 5-week-old mice from 13 CC lines (CC001, CC002, CC003, CC006, CC011, CC013, CC019, CC024, CC040, CC042, CC051, CC059, and CC061) with 1000 FFU of ZIKV (or diluent alone) by subcutaneous inoculation in the footpad. We evaluated 3 ZIKV strains (**Table 2**) with distinct virulence phenotypes in immunodeficient mice. H/PF/2013 (a 2013 human isolate from French Polynesia) causes 100% lethality in *Ifnar1*^−/−^ mice (2, 13). MR766 is the prototype ZIKV strain and has been passaged ∼150 times through suckling mouse brains, which could result in mouse-adaptive mutations (14-16). However, the MR766 variant used in these studies contains a 4 amino acid deletion ablating an N-linked glycosylation site in the viral envelope protein, resulting in a virus that is less virulent in *Ifnar1*^−/−^ mice compared to H/PF/2013 (2, 17). Dakar-MA was generated by passaging a ZIKV strain isolated from mosquitos in Senegal in the 1980s (Dakar 41525) serially in *Rag1*^−/−^ mice, resulting in a virus with enhanced pathogenesis in mice (4). Mice were weighed daily after infection, as weight loss is a sign of ZIKV disease in immunocompromised mouse models and typically precedes lethality (2). In contrast to IFN-deficient mice (and similar to WT C57Bl/6J mice) (2), none of the CC lines tested lost weight following infection with any ZIKV strain (**Figure 1**). Indeed, most lines exhibited modest weight gain over the 14 day experiment, and the rate of weight gain was no slower in infected mice than in mock-infected controls. None of the mice exhibited disease signs consistent with those observed in IFN-deficient mice (e.g. hunching, ruffled fur, paralysis, or encephalitis). CC051 and CC059 share more genetic similarities than average pairs of CC mice, so phenotypes that differ between these two lines may be especially easy to map by quantitative trait locus (QTL) analysis. Because 5-week-old CC059 mice exhibited slightly slower weight gain after ZIKV infection compared to mock-infected mice, whereas ZIKV infection had no effect on the rate of weight gain in CC051 mice, we tested ZIKV infection in weanlings of these 2 lines. We reasoned that because 3-week-old mice gain weight at a faster rate than 5-week-old mice, they may reveal phenotypes that are not evident in 5-week-old mice. However, while we found that 3-week-old mice indeed gained weight at a faster rate than 5-week-old mice, there was no difference in the rate of weight gain in mice infected with ZIKV Dakar-MA compared to mock-infected, for either CC051 or CC059 (**Figure 2**). Altogether, these observations indicate that ZIKV does not induce significant disease in any of the 13 CC lines tested.

**Table 1:** Collaborative Cross mouse lines used in this study.

| CC Line | Oas1b allele | ZIKV strains tested | Mouse ages tested | Data collected |
| --- | --- | --- | --- | --- |
| CC001 | Null | MR766<br>H/PF/2013<br>Dakar-MA | 5wk | Weight<br>Viremia<br>Neutralizing Abs |
| CC002 | Null | MR766<br>H/PF/2013 | 5wk | Weight |
| CC003 | WSB | MR766<br>H/PF/2013 | 5wk | Weight |
| CC004 | PWK | Dakar-MA | 15-21wk | Viremia |
| CC005 | Null | Dakar-MA | 15-21wk | Viremia |
| CC006 | Null | MR766<br>H/PF/2013 | 5wk | Weight |
| CC011 | Null | MR766<br>H/PF/2013 | 5wk | Weight |
| CC012 | WSB | Dakar-MA | 15-21wk | Viremia |
| CC013 | Null | MR766<br>H/PF/2013 | 5wk | Weight |
| CC019 | WSB | MR766<br>H/PF/2013 | 5wk | Weight |
| CC024 | Null | MR766<br>H/PF/2013 | 5wk | Weight |
| CC028 | PWK | Dakar-MA | 15-21wk | Viremia |
| CC040 | Null | MR766<br>H/PF/2013<br>Dakar-MA | 5wk | Weight<br>Neutralizing Abs |
| CC041 | CAST | Dakar-MA | 15-21wk | Viremia |
| CC042 | WSB | MR766<br>H/PF/2013<br>Dakar-MA | 5wk | Weight<br>Neutralizing Abs |
| CC046 | CAST | Dakar-MA | 15-21wk | Viremia |
| CC051 | Null | H/PF/2013<br>Dakar-MA | 3wk, 5wk | Weight<br>Viremia<br>Neutralizing Abs |
| CC059 | Null | MR766<br>H/PF/2013<br>Dakar-MA | 3wk, 5wk,<br>15-21wk | Weight<br>Viremia<br>Neutralizing Abs |
| CC061 | Null | MR766<br>H/PF/2013 | 5wk | Weight |
| CC068 | Null | Dakar-MA | 15-21wk | Viremia |
| CC072 | Null | Dakar-MA | 15-21wk | Viremia |

**Table 2:**
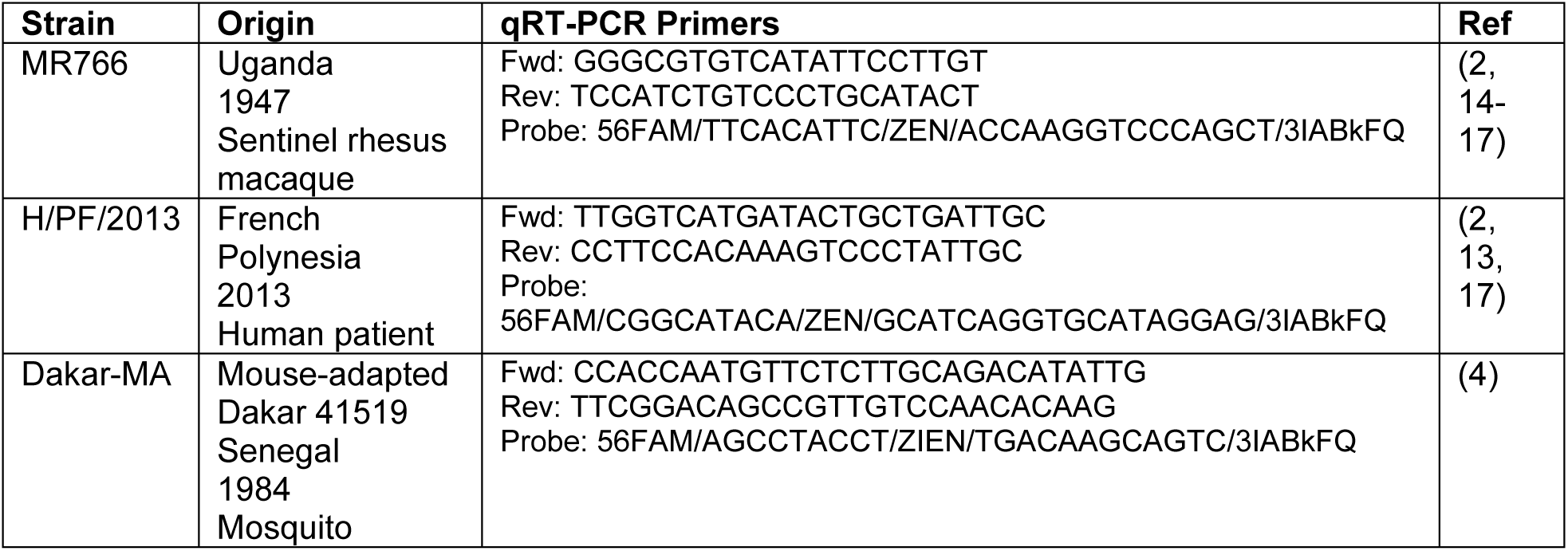
Zika virus strains used in this study.

**Figure 1:**
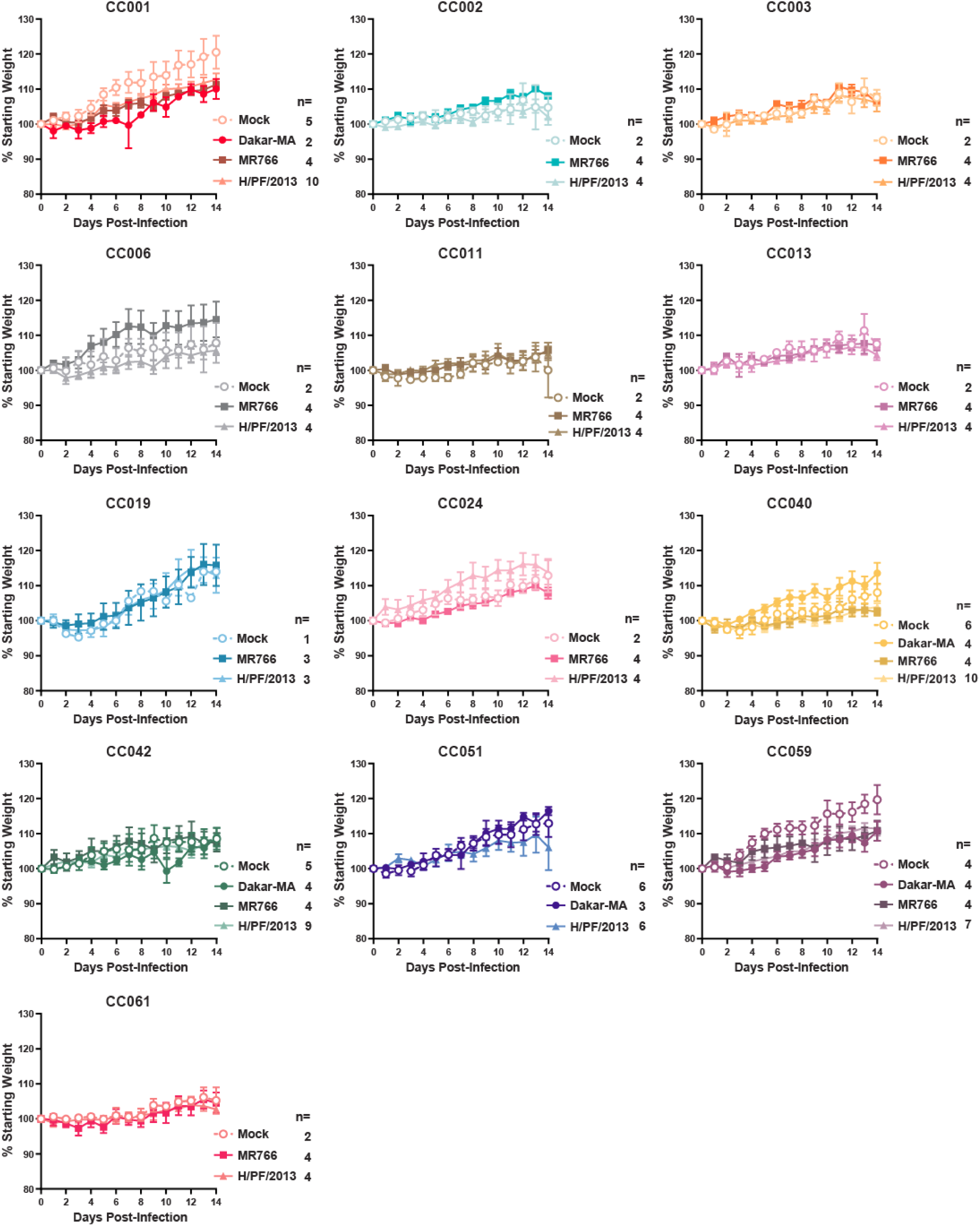
Zika virus infection does not induce weight loss in 13 Collaborative Cross lines. 5-week-old male and female mice were infected by subcutaneous footpad inoculation with ZIKV strain MR766 (1000 FFU), H/PF/2013 (1000 FFU), Dakar-MA (1 × 10^5^ FFU), or diluent (mock). Mice were weighed daily for 14 days. Weights are expressed as the percent of starting weight (day of infection). Data are shown as the mean +/− SEM of the indicated number of mice per group and are combined from 5 experiments.

**Figure 2:**
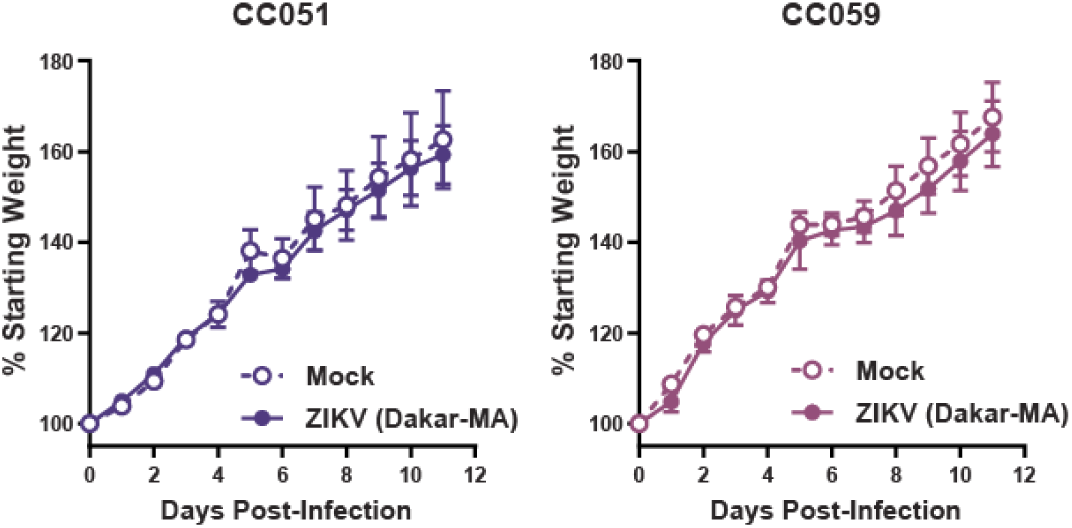
Zika virus infection does not induce weight loss in weanling CC mice. 3-week-old male and female CC051 or CC059 mice were infected by subcutaneous footpad inoculation with ZIKV strain Dakar-MA(1 × 10^5^ FFU) or diluent (mock). Mice were weighed daily for 12 days. Weights are expressed as the percent of starting weight (day of infection). Data are shown as the mean +/− SEM of 7-9 mice (CC051) or 5-6 mice (CC059) per group and are combined from 2 experiments.

### Zika virus viremia is rapidly cleared in CC mice

Although WT C57Bl/6J mice do not develop disease signs or lose weight following ZIKV infection, they do exhibit some low-level viremia which is rapidly cleared (2). We reasoned that CC lines that developed elevated or persistent viremia could provide a useful experimental system, even if the mice did not develop overt disease. To test this, we infected 5-week-old mice from 5 CC lines (CC001, CC040, CC042, CC051, and CC059) with ZIKV Dakar-MA or H/PF/2013 and measured viral RNA in serum at 1, 3, and 5 days dpi by qRT-PCR. After ZIKV Dakar-MA infection, low-level viremia (approximately 10 FFU equivalents of viral RNA per mL of serum) was detected at 1 dpi but it was greatly diminished by 3dpi (**Figure 3**). All CC001, CC042, and CC051 mice exhibited detectable viremia at 1dpi, whereas viremia was detected in only a subset of CC040 and CC059 mice. No viral RNA was detected in mice infected with ZIKV H/PF/2013, though viremia was evaluated only at 3 dpi and only in CC042, CC051, and CC059 mice. We next evaluated ZIKV viremia in adult CC mice (15-21 weeks old) infected with ZIKV Dakar-MA. Similar to 5-week-old mice, adult mice exhibited approximately 10 FFU equivalents of viral RNA per mL of serum at 1dpi, which diminished by 3dpi and was undetectable by 5dpi (**Figure 4**). Altogether, these results show that while some ZIKV replication does occur in CC mice, the virus is cleared rapidly, consistent with the lack of disease signs observed in CC mice. Importantly, we did not observe robust differences in viremia between CC lines, indicating that this is not a suitable phenotype for QTL mapping studies.

**Figure 3:**
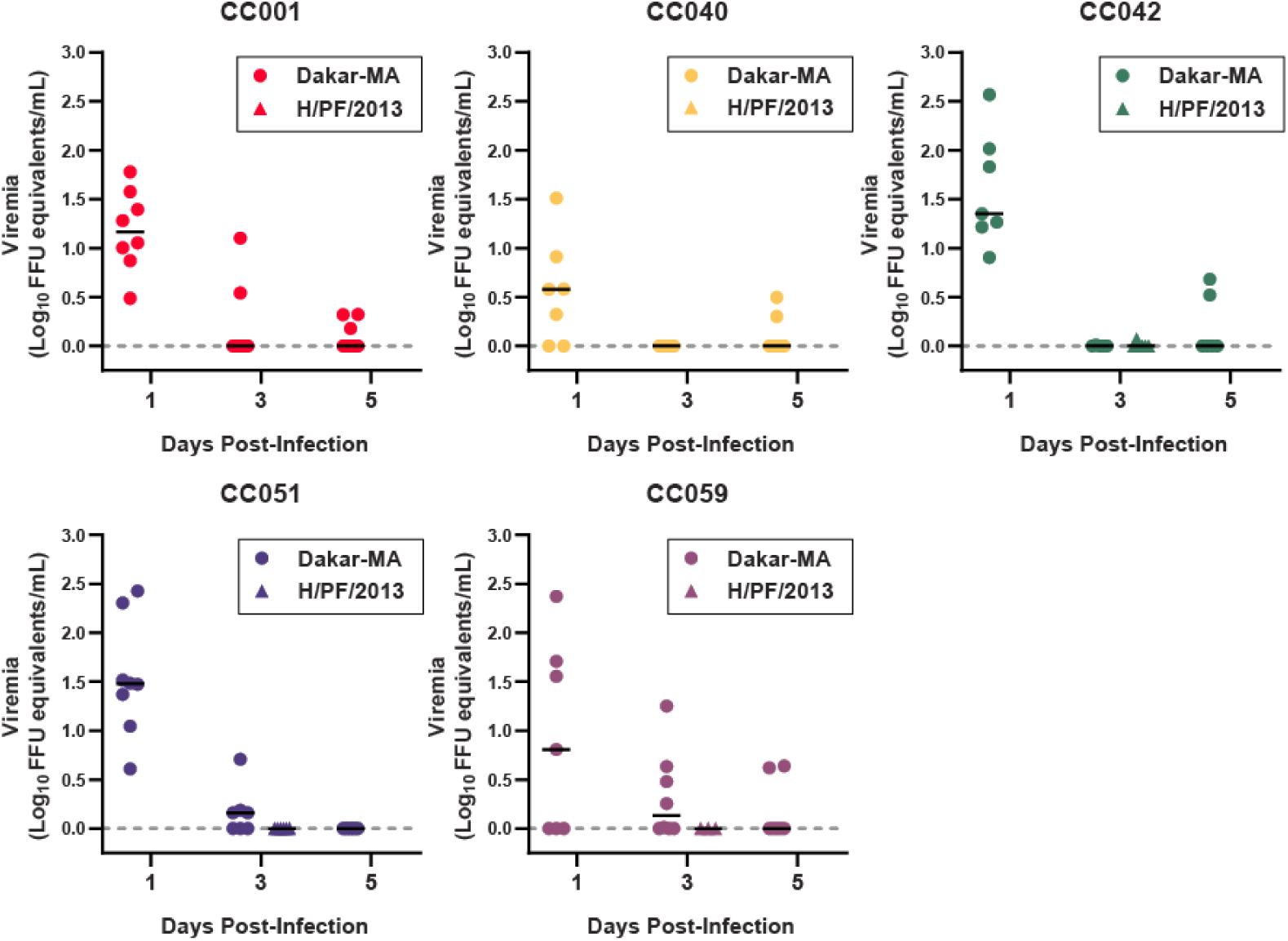
Zika virus viremia is rapidly cleared in young CC mice. 5 week-old male and female mice of the indicated CC lines were infected by subcutaneous footpad inoculation with ZIKV strain Dakar-MA (1 × 10^5^ FFU) or H/PF/2013 (1000 FFU). Blood was sampled at 1, 3, and 5 days post-infection and viral RNA in serum was measured by qRT-PCR. Dotted line indicates limit of detection. Each data point corresponds to one mouse and data are combined from 3 experiments; bar indicates median.

**Figure 4:**
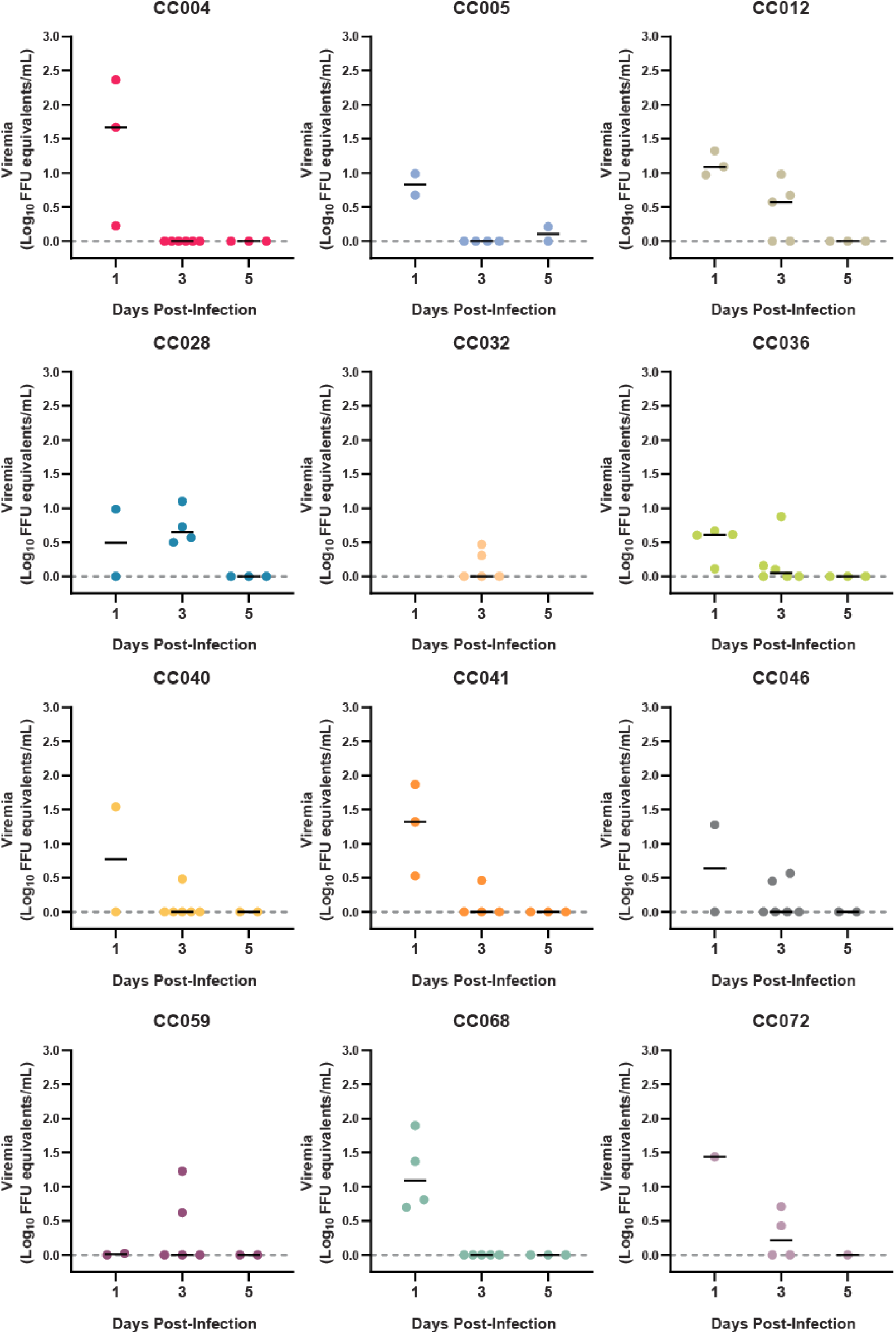
Zika virus viremia is rapidly cleared in adult CC mice. 15-21 week-old male mice of the indicated CC lines were infected by subcutaneous footpad inoculation with ZIKV strain Dakar-MA (1 × 10^5^ FFU). Blood was sampled at 1, 3, and 5 days post-infection and viral RNA in serum was measured by qRT-PCR. Dotted line indicates limit of detection. Each data point corresponds to one mouse and data are combined from 5 experiments; bar indicates median.

### Zika virus infection induces neutralizing antibodies in Collaborative Cross mice

Since ZIKV produced only transient low-level viremia in CC mice, we tested whether this was sufficient to elicit the production of neutralizing antibodies. We collected serum at 14dpi from 5 CC lines (CC001, CC040, CC042, CC051, CC059) infected with ZIKV H/PF/2013 at 5 weeks of age. Heat-inactivated serum was tested for neutralizing activity against ZIKV H/PF/2013 and MR766 (**Figure 5**). Consistent with low-level viremia, neutralizing antibody titers were low (median FRNT50 titers <10), with the exception of CC042 which exhibited FRNT50 titers of ∼100. CC042, CC001, and CC051 exhibited similar viral loads at 1dpi, indicating that viremia is not the sole determinant of neutralizing antibody responses. Consistent with our prior studies of human antibody responses to ZIKV (23), MR766 was more readily neutralized (higher FRNT50) than H/PF/2013.

**Figure 5:**
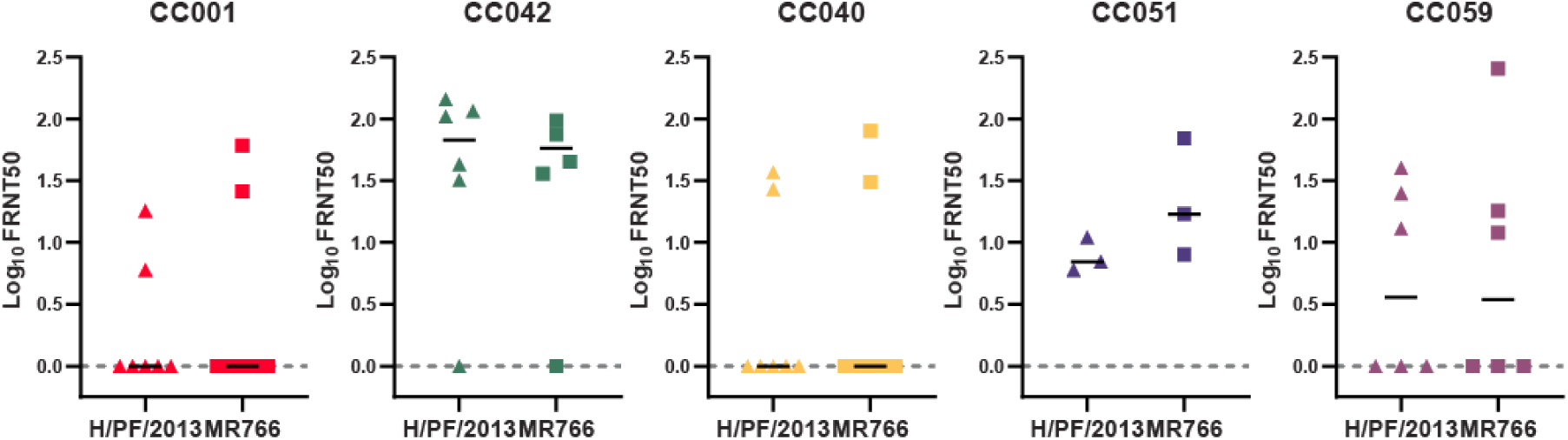
Zika virus infection induces neutralizing antibodies in Collaborative Cross mice. 5-week-old male and female mice were infected by subcutaneous footpad inoculation with ZIKV strain H/PF/2013 (1000 FFU) and serum was collected at 14 days post-infection. Heat-inactivated serum was tested for neutralizing activity against ZIKV strains H/PF/2013 and MR766 by focus-reduction neutralization test on Vero cells. FRNT50 values indicate the serum dilution at which infection was inhibited by 50%. Dotted line indicates limit of detection. Each data point corresponds to one mouse; bar indicates median,

Taken together, our results are consistent with other studies demonstrating poor ZIKV infection in IFN-intact mice and suggest that the tested CC lines do not include polymorphic host genes that greatly increase susceptibility to ZIKV infection.

## Discussion

Prior to the emergence of ZIKV in Latin America, few studies had evaluated ZIKV pathogenesis in animals (24-26). However, the explosive growth of ZIKV research since 2015 has spurred the development of new animal models, revealing pathogenic mechanisms and providing systems for evaluating therapeutic interventions. The most significant models of ZIKV infection are non-human primates and mice (7), but ZIKV disease also has been studied in guinea pigs, hamsters, tree shrews, and swine (27-35). Mice offer significant advantages in terms of cost, scale, speed, and genetic tractability, enabling mechanistic studies that are not feasible in non-human primate models. A significant disadvantage of mouse models is that ZIKV replicates poorly in immunocompetent mice and thus does not exhibit the tissue tropism and disease phenotypes characteristic of human infection. Mice lacking type I IFN production or responses succumb to ZIKV infection and exhibit broad tissue tropism, implying that the innate antiviral response is a key barrier to ZIKV replication and pathogenesis in mice (2, 36). Mice lacking the IFN-αβ receptor (alone, or also lacking the IFN-γ receptor) have provided useful models for studying ZIKV disease phenotypes including congenital infection, sexual transmission, and ocular infection, as well as for evaluating candidate vaccines and antivirals (37-43). The restriction of ZIKV replication in mice results in part from an inability of ZIKV NS5 to degrade murine STAT2 and therefore ZIKV cannot inhibit IFN signaling in mice by the same mechanism used in human cells (5, 6). However, NS5-mediated antagonism restricts antiviral signaling only in infected cells whereas these responses are sustained in uninfected cells responding to paracrine IFN signaling. Thus, *Stat2*^−/−^ mice succumb to ZIKV infection, but transgenic mice expressing human STAT2 only exhibit disease in the context of ZIKV strains with additional mouse-adaptive mutations (4, 44). In addition, the ZIKV NS2B-NS3 protease targets human but not murine STING, resulting in sustained IFN production and diminished viral replication in mouse cells (3). However, mice lacking STING were not more susceptible to ZIKV than wild-type mice (2, 3), implying that STING is a contributing but not dominant factor restricting ZIKV infection in mice.

Flavivirus resistance is one of the earliest examples of a genetic determinant of pathogen susceptibility defined in mice. Over 80 years ago, flavivirus resistance was shown to be inherited in a single gene autosomal dominant manner (22, 45, 46), and more than 15 years ago the resistance phenotype was mapped to the *Oas1b* gene (47, 48). More recent studies used F1 hybrids of CC mice to define genetic determinants of WNV pathogenesis and found that the QTL with the largest effect mapped to *Oas1b* (12, 49). The antiviral activity of Oas1b restricts all flaviviruses (>13 viruses tested) and appears to act exclusively against flaviviruses. Other members of the Oas gene family restrict viral replication by sensing dsRNA and synthesizing the second messenger 2’-5’ oligoadenylate, which activates the cytoplasmic RNA degrading enzyme RNaseL. However, Oas1b is not catalytically active and does not activate RNaseL (50). The flavivirus restricting activity of Oas1b is cell intrinsic and restricts viral replication in a post-entry manner, but its mechanism is unknown (50, 51). OAS orthologs in other species also restrict viral replication (52, 53), and *OAS1* polymorphisms are associated with WNV disease in humans (54, 55) and horses (56). However, humans do not have a functional equivalent of murine *Oas1b* with RNaseL-independent inhibitory activity. Thus, identifying non-Oas1b host factors that modulate flavivirus susceptibility in mice may provide insight into mechanisms that control flavivirus disease in humans.

There is a need for improved mouse models of ZIKV infection to advance studies of ZIKV pathogenic mechanisms and to evaluate candidate vaccines and therapeutics. Current ZIKV mouse models typically rely on mice that lack type I IFN signaling which limits their utility for studying immune mechanisms that control ZIKV replication and disease. In particular, IFN signaling plays a key role in restricting flavivirus tropism and IFN-deficient mice typically develop disseminated ZIKV infection that does not recapitulate the tissue tropism observed in humans. CC mice have provided useful experimental systems for other pathogenic viruses whose disease phenotypes are not well-modeled in conventional laboratory mouse lines, including Ebola virus, SARS coronavirus, and chikungunya virus (10, 11), because they exhibit unique phenotypes that are not necessarily predicted from the 8 founder lines. Thus, we sought to determine whether CC mice might provide a useful model system for ZIKV. We found that, although ZIKV viremia was detectable at 1dpi, the amount of viral RNA detected in serum was very low and cleared rapidly, and infected mice exhibited no signs of disease including weight loss. Based on these results, the tested CC lines are unlikely to be useful as models of ZIKV disease or as tools to map host factors that control ZIKV pathogenesis. This was true even of CC lines carrying non-functional alleles of *Oas1b*, which would be expected to be the most susceptible to flavivirus infection (12).

It is possible that additional approaches could reveal experimentally useful ZIKV phenotypes in CC mice. These could include administering virus by an intravenous, rather than subcutaneous route, or infecting with a higher dose of virus, although the 1×10^5^ FFU dose we used for infections with ZIKV Dakar-MA is quite high. Another approach would be administering an IFNAR-blocking monoclonal antibody (MAR1-5A3) prior to ZIKV infection. In C57Bl/6J mice, MAR1-5A3 treatment results in elevated viremia and relevant clinical phenotypes such as transplacental transmission, though it is not sufficient to produce weight loss or lethality. Furthermore, our study tested only a subset of CC lines; additional lines not tested here may reveal more robust pathogenesis phenotypes. The overall lack of disease in CC mice highlights the species-specific restriction factors that limit ZIKV infection in mice. CC mice remain a valuable system for studying host factors that control flavivirus pathogenesis, but are likely to be more revealing for flaviviruses that are able to cause more robust infection in mice than ZIKV.

## Acknowledgements

This work was supported by U19-AI100625-04S1 (M.T.H. and R.S.B.), a Pilot Project from the UNC Systems Genomics program (H.M.L.), and startup funds from UNC Department of Microbiology and Immunology and the Lineberger Comprehensive Cancer Center (H.M.L.)

